# A high-content screen identifies the vulnerability of MYC-overexpressing cells to dimethylfasudil

**DOI:** 10.1101/801134

**Authors:** Jing Zhang, Shenqiu Zhang, Qiong Shi, Thaddeus D. Allen, Fengming You, Dun Yang

**Author notes:** Corresponding authors (DY), (FY).

## Abstract

A synthetic lethal effect arises when a cancer-associated change introduces a unique vulnerability to cancer cells that makes them unusually susceptible to a drug’s inhibitory activity. The synthetic lethal approach is attractive because it enables targeting of cancers harboring specific genomic alterations, the products of which may have proven refractory to direct targeting. An example is cancer driven by overexpression of MYC. Here, we conducted a high-content screen for compounds that are synthetic lethal to elevated MYC using a small-molecule library to identify compounds that are closely related to, or are themselves, regulatory-approved drugs. The screen identified dimethylfasudil, a potent and reversible inhibitor of Rho-associated kinases ROCK1 and ROCK2. Close analogs of dimethylfasudil are used clinically to treat neurologic and cardiovascular disorders. The synthetic lethal interaction was conserved in rodent and human cell lines and could be observed with activation of either MYC or its paralog MYCN. The synthetic lethality seems specific to MYC overexpressing cells as it could not be substituted by a variety of oncogenic manipulations and synthetic lethality was diminished by RNAi-mediated depletion of MYC in human cancer cell lines. Collectively, these data support investigation of the use of dimethylfasudil as a drug that is synthetic lethal for malignancies that specifically overexpress MYC.

**Significance Statement:** Synthetic lethal targeting of tumors overexpressing MYC holds promise for attacking aggressive malignancies. Here we describe a synthetic lethal interaction between dimethylfasudil and overexpression of MYC. Uniquely, this novel synthetic lethal interaction points toward an opportunity for synthetic lethality with a molecule likely to harbor favorable drug-like properties that enable systemic use.

## Introduction

At least one of the proteins encoded by *MYC*, *MYCN* and *L-MYC*, paralogous members of the MYC family, are deregulated in over 50% of human cancers. This can be due to mutation, translocation, amplification, or alterations in upstream regulators [1–3]. Overexpression of MYC is directly transforming [4] and several studies have shown that diminishing MYC *in vivo* can elicit tumor regression, suggesting that direct therapeutic targeting of MYC will be a way to attack human malignancies [5–7].

Major challenges exist to developing a drug that directly inhibits MYC. As a transcription factor, MYC functions in the nucleus. Any drug that inhibits MYC may have to gain access to the nuclear compartment, something currently only feasible for small molecule drugs. However, unlike kinases, MYC lacks a specific active site for such a small molecule inhibitor to interact. In addition, tumor cell heterogeneity can contribute to the emergence of clones that have acquired additional mutations that allow MYC-independent growth [8]. This means that not all cancer cells with abundant MYC will be absolutely dependent on MYC for proliferation and survival.

One strategy that might overcome all these obstacles is to exploit cellular dependencies uniquely induced by MYC in cancer cells. Such synthetic lethality between overexpression of MYC and targeting of other cellular pathways and proteins has been described with the activation of the DR5 death receptor pathway [9], depletion of the non-essential amino acid glutamine [10], depletion of the core spliceosome component, BUD31 [11], depletion of AMPK-related kinase 5 [12] and pharmacological inhibition of CDK1 [13], PIM1 [14] and Aurora kinase B (AURKB) [15]. Unfortunately, the clinical potential for AURKB inhibitors has been hampered by patient toxicity. This may also limit other synthetic lethal approaches that have not yet advanced to the clinic.

To bypass this possibility, we conducted a phenotypical screen using a small-molecule library of already clinically used drugs and their close analogs for the ability to elicit cell-division defects specifically in MYC overexpressing cells. The high-content screen positively identified a ROCK kinase inhibitor dimethylfasudil (diMF), but not its analogs fasudil and ripasudil. These analogs are themselves ROCK inhibitors so synthetic lethality is likely mediated by an as yet to be defined target unique to diMF. We utilized both model cell lines and human cancer cell lines to confirm the MYC-diMF synthetic lethal interaction. It was conserved in rodent and human cells and was, in all cases, associated with induction of classical apoptotic features and accumulation of polyploid cells.

Fasudil and ripasudil are both used clinically, with fasudil being used as a systemic treatment. It is likely that diMF may also possess drug-like properties such as a favorable pharmacokinetic and adverse event profile that would enable clinical use. Our findings suggest a new avenue for developing a potent and non-toxic therapeutic intervention for MYC overexpressing tumors using diMF.

## Materials and Methods

### Cell lines and culture conditions

Human cancer cell lines that were purchased from the American Type Culture Collection (ATCC) include Hela (ATCC^®^ CCL-2™), NCI-H841 (ATCC^®^ CRL-5845™), DU-145 (ATCC^®^ HTB-81™) and Calu-6 (ATCC^®^ HTB-56™). Cells were grown for three passages and frozen as cell stock. Prior to each experiment, a stock vial was thawed and serially passaged for no more than five generations. As previously described [13], RPE-NEO and RPE-MYC cells were generated by transfecting hTERT-immortalized human primary retinal pigment epithelial cells (RPE-hTERT) with a vector expressing the Neomycin resistance gene (*Neo*) alone or alongside the *MYC* oncogene, respectively, followed by selection with 800 μg/ml G418/Geneticin. RPE-MYC cells were transfected with a construct harboring an *H2B-GFP* fusion gene and selected with 1 μg/ml of Puromycin to generate RPE-MYC^H2B-GFP^ cells for high-content screening. The mammalian *H2B-GFP* expression construct was a gift from Dr. Andrei Goga of University of California, San Francisco. Construction of derivatives of Rat1A cells and verification of ectopic expression of oncogenic proteins has been described [13, 16]. Briefly, Rat1A cells were infected with retrovirus expressing a variety of cancer genes. Transduction efficiency was judged using GFP expression. Rat1A/MycN^ER^, IMR-90/Myc^ER^, HA1E/Myc^ER^, and HA1E/MycN^ER^ were generated by infection of parental cells with viral particles produced using Phoenix packaging cells (ATCC^®^ CRL-3213™) followed by Puromycin selection (1 μg/ml) [10, 17]. All mammalian cells were cultured in DMEM supplemented with 10% fetal bovine serum and antibiotics at 5% CO2 in a humidified incubator. To ensure that cell lines were free of mycoplasma contamination, a Universal Mycoplasma Detection Kit (ATCC^®^ 30-1012K™) was used to assay for the mycoplasma 16S rRNA-coding region after experimentation.

### Cell viability assays

Cell viability was determined by the Trypan blue exclusion assay, as previously described [15], or alternatively through immunofluorescent staining for various apoptotic markers. Viability was expressed as the percentage of dead cells in the population at an indicated time-point following initiation of drug treatment. For each assay, data is presented from three independent experiments that were performed with sequentially passaged cells, and each experiment was done in triplicate (n=9).

### Small molecules

The FDA-approved drug panel of 2576 drugs (Cat. # L1300) was purchased from Selleck Chemicals. An additional two hundred analogs of various FDA-approved drugs were synthesized in-house and were included in our high-content screening panel. The following small molecules were from commercial sources and used at the final concentrations indicated in the figures or their legends: VX-680 (Cat. # S1048) was purchased from Selleck Chemicals; paclitaxel (Cat. # P-9600), etoposide (Cat. # E-4488), staurosporine (Cat. # S-9300), and doxorubicin (Cat. # D-4000) were obtained from LC Laboratories; nocodazole (Cat. # M1404), cycloheximide (Cat. # C-7698), cisplatin (Cat. # C2210000), and monastrol (Cat. # M8515) were from Sigma-Aldrich; Zeocin (Cat. # R25001), G418/Geneticin (Cat. # 10131035), 0.4% trypan blue solution (Cat. # 15250061), and puromycin dihydrochloride (Cat. # A1113802) were purchased from Thermo-Fisher Scientific; Eg5 inhibitor II (Cat. # BP-30219) was obtained from BroadPharm; diMF (Cat. # sc-203592) and dihydrocytochalasin B (DCB) (Cat. # sc-202579) were acquired from Santa Cruz Biotechnology.

### High-content drug screening for MYC-synthetic lethal agents

RPE-MYC^H2B-GFP^ cells were cultured in 150 mm petri dishes and were cryo-preserved in large amounts before use. Cells with the same passage number were thawed and seeded to batches of 96-well microtiter plates with 2,000 cells per well as the starting cell number for high-content screening. Treatment with small molecules was initiated 24 hours after seeding at a final concentration of 10 μM in 0.1% DMSO. Cells were imaged 24 and 48 hours after initiation of drug treatment to monitor H2B-GFP-labelled DNA using an IN Cell Analyzer 2000 (GE). Compounds that triggered mitotic arrest at 24 hours and polyploidy and apoptosis at 48 hours were scored as positive and were subjected to further validation.

### RNA interference

Cells were cultured in a 6-well plate and allowed to adhere overnight before transfection with either a negative control siRNA, or *MYC* siRNA at a concentration of 50 nM using Lipofectamine 2000 (Invitrogen) according to the manufacturer’s instructions. The sequences for the *MYC* siRNA and control siRNA were previously described [18]. The transfection efficiency was more than 95% as judged by expression of a co-transfected pEGFP-N1 construct (Clontech).

### Immunofluorescent microscopy

Immunofluorescent staining experiments were performed as previously described [15]. Cells were cultured on coverslips in a 6-well plate, fixed with either 4% paraformaldehyde or 4% paraformaldehyde followed with methanol treatment, and then permeabilized with 0.3% Triton X-100. Rabbit IgG antibodies for active caspase 3 (Cat. # 9661S), active caspase 9 (Cat. # 20750S), and cleaved PARP (Cat.# 5625S) were from Cell Signaling Technology and used at 1:100 dilution for each. Primary antibodies were detected with Rhodamine (TRITC)-conjugated AffiniPure Donkey Anti-Rabbit IgG (H+L) (Jackson ImmunoResearch, Cat. # 711-025-152) at 1:500 dilution. After immunostaining, we mounted cells on microscope slides with 4’,6’-diamidino-2-phenylindole (DAPI)-containing Vectashield mounting solution (Vector Laboratories, Cat. # H1500). For determination of mitochondrial membrane potential, cells were cultured on cover slips and exposed for 30 min. at 37 °C with 40 nM Mitotracker® Red CMXRos (Thermo Fisher, Cat. # M7512), a dye that accumulates in mitochondria as a function of the mitochondrial membrane potential. After fixation in 4% paraformaldehyde, Mitotracker® Red-stained cells were mounted on microscope slides with DAPI-containing Vectashield. For fluorescence detection, we used either a Zeiss LSM510 confocal laser microscope with a 63X/1.4 N.A. oil objective or an EVOS FL Auto microscope (Thermo Fisher).

### Western blot analysis

Whole cell extracts were prepared by treating cells for 15 min. at 4 °C with a lysis buffer [50 mM Tris (pH 7.5), 200 mM NaCL, 0.1% SDS, 1% Triton X-100, 0.1 mM DTT, and 0.5 mM EGTA] supplemented with BD BaculoGold Protease Inhibitor Cocktail (BD Biosciences, Cat. # 554779). The extracts were centrifuged at 8,000 × g for 10 min. to clear insoluble material. The protein concentration in the supernatant was determined using a Bio-Rad Protein Assay (Bio-Rad, Cat. # 5000006). Lysates containing 50μg of protein were resolved on NuPAGE (4–12%) Bis-Tris gels (Thermo Fisher, Cat. # NP0322PK2) and transferred to nitrocellulose membranes (Bio-Rad, Cat. # 1620150). Equal transfer of protein in each lane was assessed by staining of total protein on the membrane with solution containing 0.1% Ponceau S (w/v) and 5.0% Acetic Acid (w/v) (Sigma, Cat. # P7170-1L). Membranes were then blocked with 5% nonfat milk in PBS buffer for 1 hour before incubating overnight at 4 °C with primary antibody diluted 1:1,000 in blocking buffer. Rabbit antibodies against MYC (Cat. # ab32072) and Actin (Cat. # ab179467) were from Abcam and used at 1:1000 dilution. Horseradish peroxidase-conjugated anti-rabbit immunoglobulins (Santa Cruz Biotechnology, Cat. # sc-2357) were used at 1:5000 dilution. Western blots were visualized with the SuperSignal West Femto ECL detection kit (Thermo Fisher, Cat. # 34095).

### Blinding of experimental screen and statistical analysis

To reduce observer’s bias, we blinded the screening experiments presented in Figures 1, 5 and 6. Both the experimenter and statistician were blinded when possible. We randomized the identity of inhibitors with the RANDBETWEEN function in Excel and relabeled chemical inhibitors accordingly. Although, chemical identity was blinded, the identity of cell lines was not always blinded to the experimental biologists, because they can distinguish morphological differences among different cell lines. They completed the experiment, and collected data which was statistically analyzed. The identity of cell lines and inhibitors was revealed after completion of statistical analysis.

**Figure 1:**
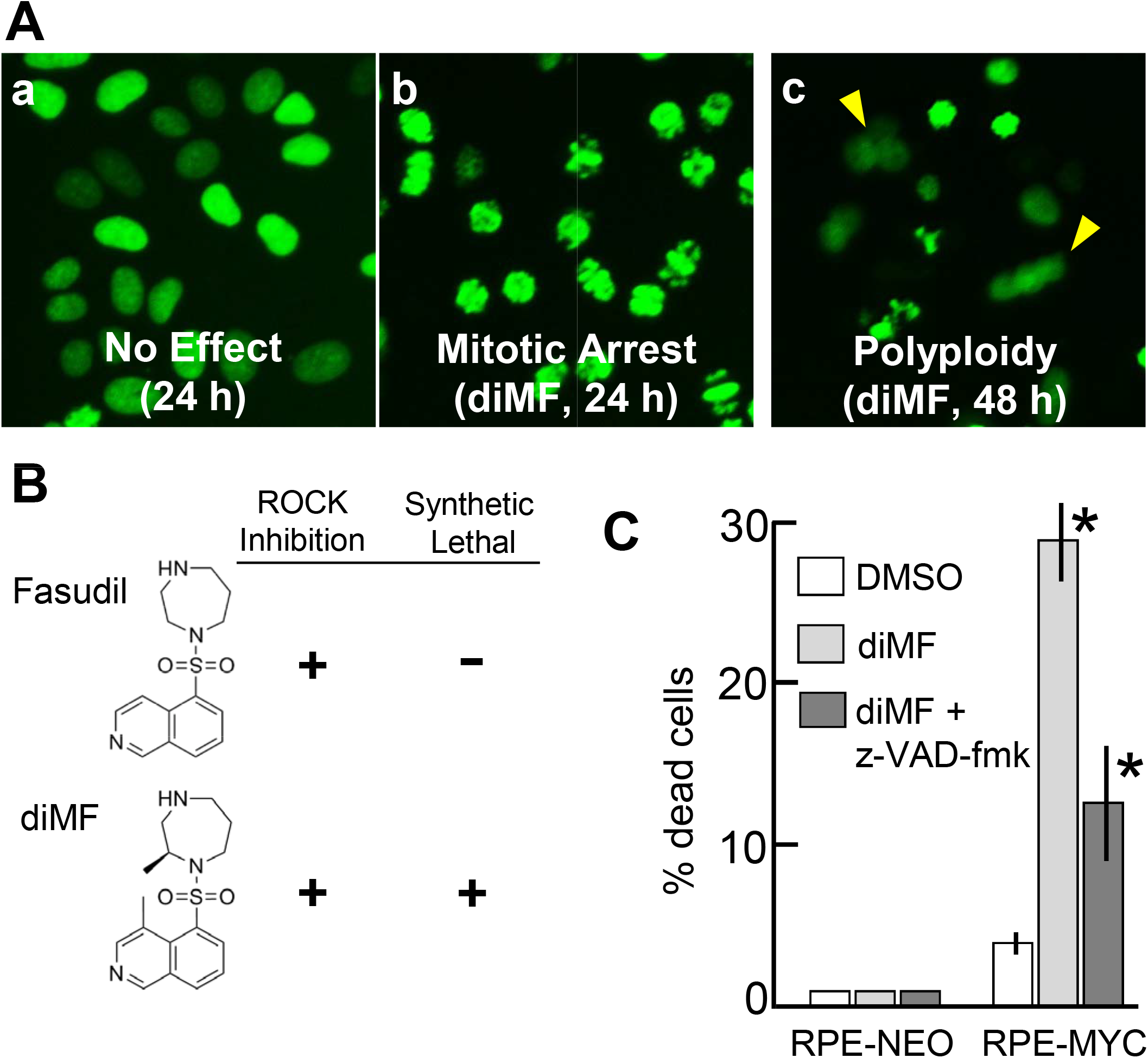
diMF elicits MYC-dependent cytotoxicity. Representative images **(A)** of RPE-MYC^H2B-GFP^ cell phenotypes observed with high-content screening with a small molecule library. Shown are cells treated with **(a)** DMSO as a negative control, **(b)** cells mitotically arrested after 24 hours and **(c)** polyploid cells (arrowheads) that formed after 48 hours of diMF (10 μM). **(B)** Comparison of the chemical structure and functional activity of fasudil and diMF. **(C)** Partial inhibition of MYC-diMF synthetic lethality by the caspase inhibitor z-VAD-fmk. RPE-NEO and RPE-MYC cells were treated for 72 hours with 0.1% DMSO (□), 6 μM diMF (■), or 6 μM diMF + 100 μM z-VAD-fmk (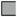). Cell viability was determined by the trypan blue exclusion assay (n=9). Error bars denote standard deviation. Statistical analysis was with one-way AVOVA followed by Dunnett’s test. The symbol * indicates *p<0.01* when compared with the DMSO control group.

Statistical analyses were performed with R software. *F* statistics derived from one-way ANOVA were computed using the aov function from the stats library. The *F* statistics tested the null hypothesis that all group means were equal and were performed at 5% significance level. When null hypothesis of the *F* test was rejected, Dunnett’s tests were performed to identify the group means that were statistically different from the control group. Dunnett’s two-sided tests were computed using the DunnettTest function from the DescTools library and were used to compare multiple treatment groups as in Figures 1C, 3 and 5. The two-tailed, unpaired t-tests in Figures 2C, 4 and 6 were computed using the t.test function from the stats library. Each data set represents collective data points from three independent experiments, with three replicates in each experiment (n=9 observations/data point). Error bars represent one standard deviation from the mean. The number of experiments and sample sizes were determined before the collection of any data. In all figures, * indicates significance at a 1% level. The F test and Dunnett Test assume normally distributed data. We tested normality of the data with the Shapiro-Wilk’s test and failed to reject the null hypothesis of normality in all cases.

**Figure 2:**
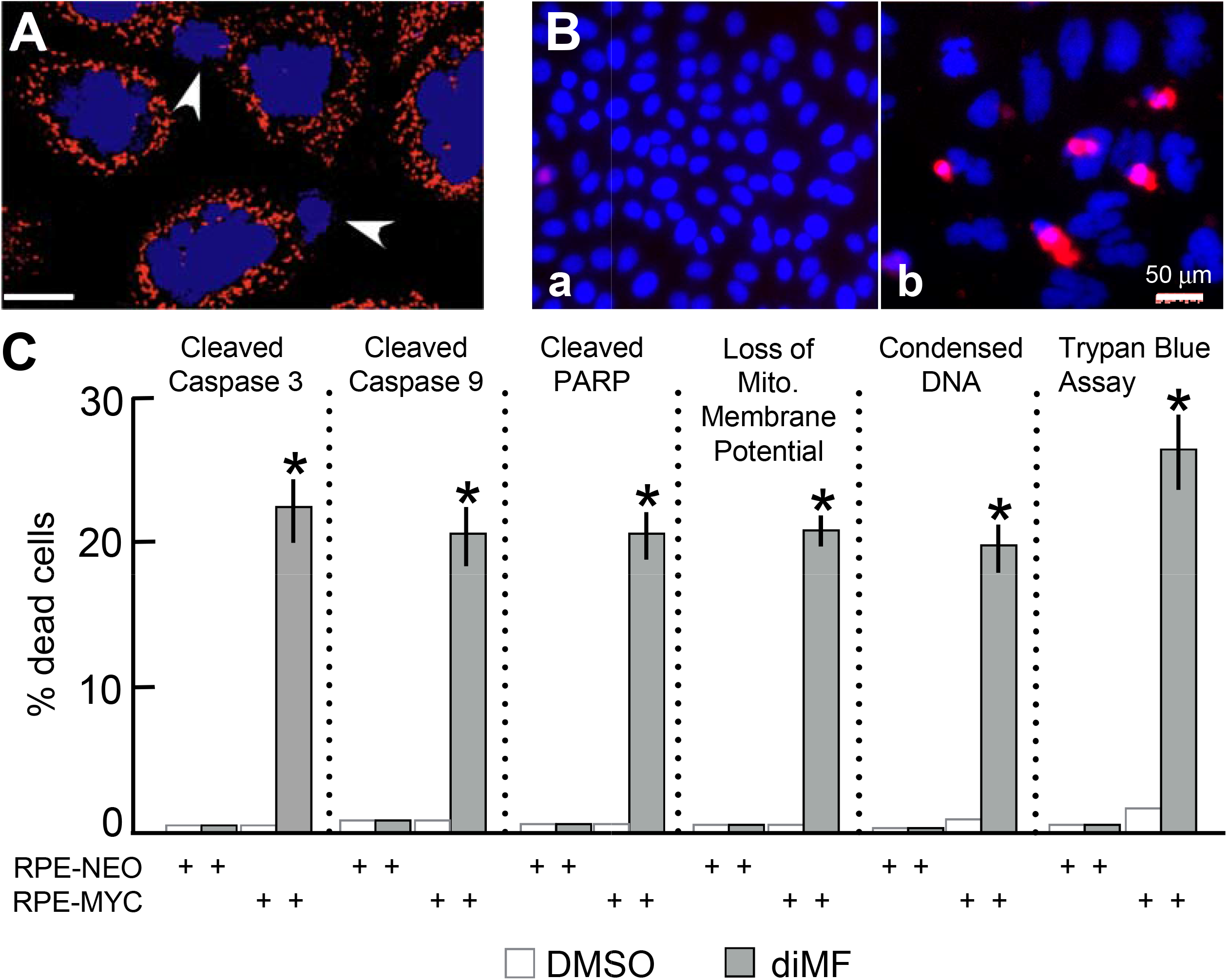
MYC-diMF engages the intrinsic apoptotic cascade for synthetic lethality. **(A)** Early apoptotic nuclei in RPE-MYC cells exposed to 6 μM diMF for 48 hours and then incubated for 30 minutes with 40 nM Mitotracker™ Red, a dye that accumulates in mitochondria and for which fluorescence intensity depends on the mitochondrial membrane potential, ΔΨm. Cells were stained for DNA with DAPI (blue) and image captured a confocal laser microscope (arrowheads indicate apoptotic nuclei). Scale bar, 10 μm. **(B)** Immunofluorescent staining for activated (cleaved) caspase 9 in RPE-MYC cells treated with 0.1% DMSO **(a)** or 6 μM diMF **(b)** for 3 days. Cells were stained for DNA with DAPI (blue). Micro-images were taken with an EVOS FL Auto microscope. Scale bar, 50 μm. **(C)** Percentage of cells positive in the indicated assays. RPE-MYC and RPE-NEO cells were treated with 0.1% DMSO (□) or 6 μM diMF (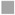) for 3 days and then subjected to the indicated viability assays. The experiment was done in triplicate so that n=9 for each data point. Error bars represent standard deviation. The symbol * indicates *p <0.01* when diMF treatment was compared with DMSO treatment (unpaired, 2-tailed t-test).

**Figure 3:**
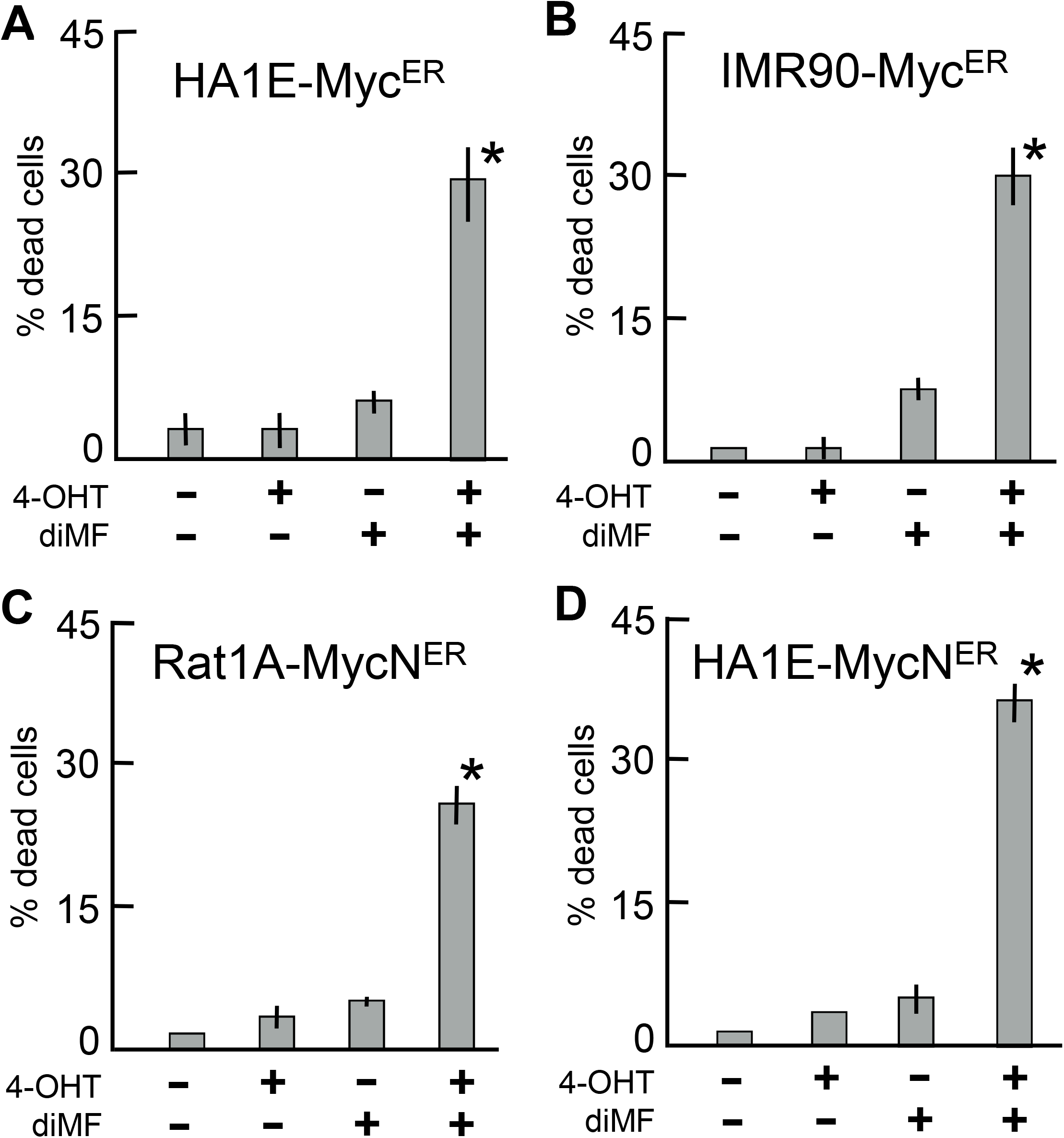
Synergistic induction of cell death by diMF in combination with acute activation of either MYC or MYCN. MYC^ER^ expressing HA1E **(A)** or IMR90 **(B)** cells and MYCN^ER^ expressing Rat1A **(C)** or HA1E **(D)** cells were incubated with or without 6 μM diMF and/or 200 nM 4-OHT for 3 days. Cell viability, was determined by the trypan blue exclusion assay. Each column represents the average of three independent experiments with three replicates in each (n=9). Error bars represent one standard deviation. Statistical analysis was by one-way ANOVA with *p-values* calculated using the Dunnett’s test. The symbol * indicates *p<0.01* when the indicated double treatment was compared with any of the no treatment or individual treatment groups.

**Figure 4:**
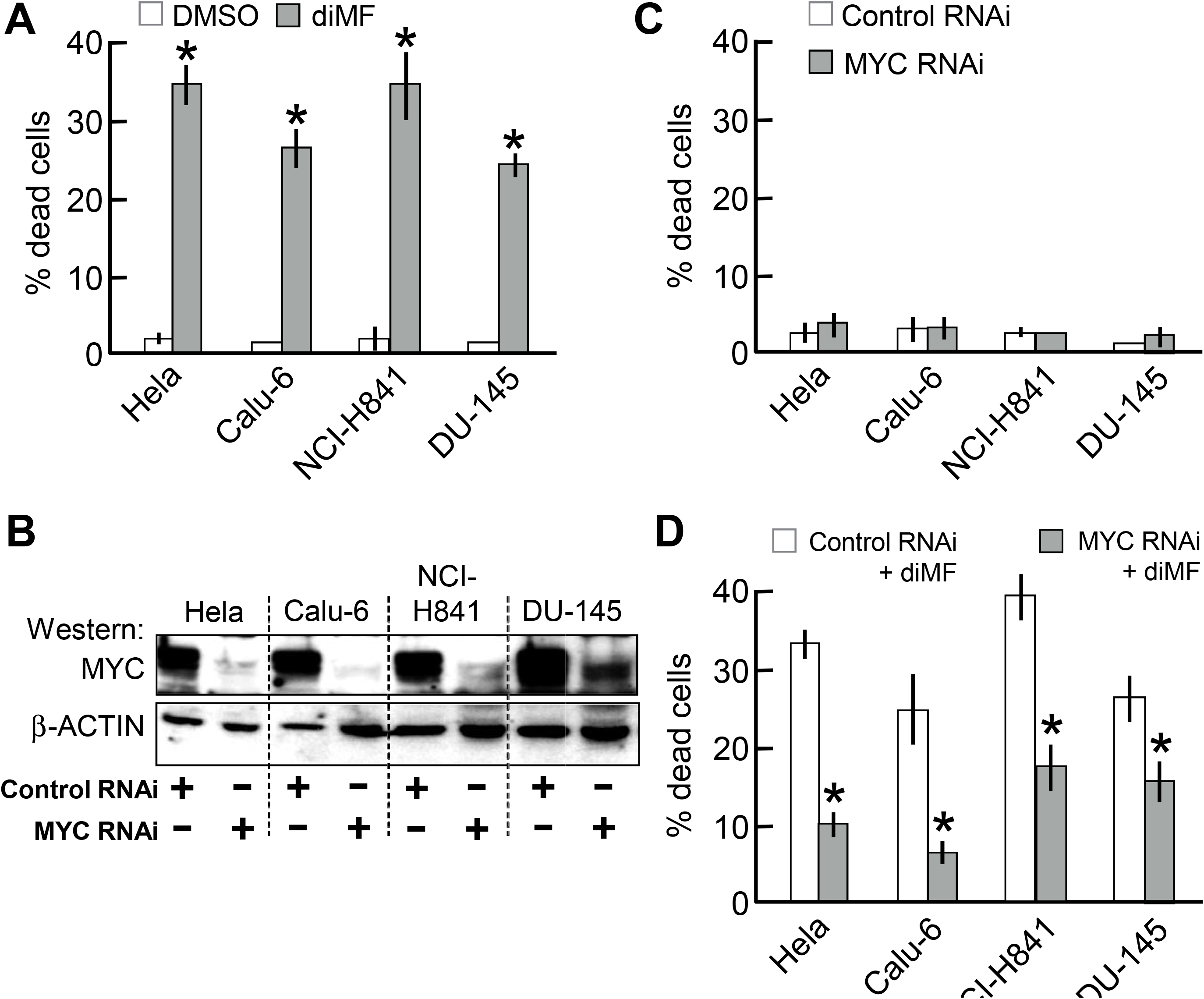
MYC is required in cancer cell lines for cytotoxic killing by diMF. **(A)** Lethal effect of diMF on a variety human cancer cells that overexpress MYC. Cells were treated with 0.1% DMSO (□) or 6 μM diMF (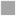) for 3 days. Cell viability was assayed by Trypan blue exclusion assay. **(B)** Depletion of MYC in human cancer cell lines by RNAi. The indicated human cancer cell lines were transfected with either control siRNA (lanes 1, 3, 5, and 7) or *MYC* siRNA (lanes 2, 4, 6, and 8). Extracts were prepared 3 days after siRNA transfection and Western analysis used to examine levels of MYC and β-ACTIN protein. **(C)** Depletion of MYC alone fails to elicit cell death. The indicated cancer cell lines were transfected with either control siRNA or *MYC* siRNA. Cell viability was determined by the trypan blue exclusion assay 4 days after transfection. **(D)** Suppression of diMF cytoxicity by RNAi-depletion of MYC. The indicated cancer cell lines were transfected with either control siRNA or *MYC* siRNA and then exposed to 6 μM diMF starting 24-hours after transfection. Cell viability was determined by the trypan blue exclusion assay at day 4 after transfection. For data in **A**, **C**, and **D**, each data point represents the average of 3 independent experiments with data collected in triplicate (n= 9). Error bars represent the standard deviation. The pairwise comparisons were done using two tailed, unpaired *t*-tests. The symbol * indicates *p<0.01* compared to DMSO for **A** and compared to control RNAi groups in **C** and **D**.

**Figure 5:**
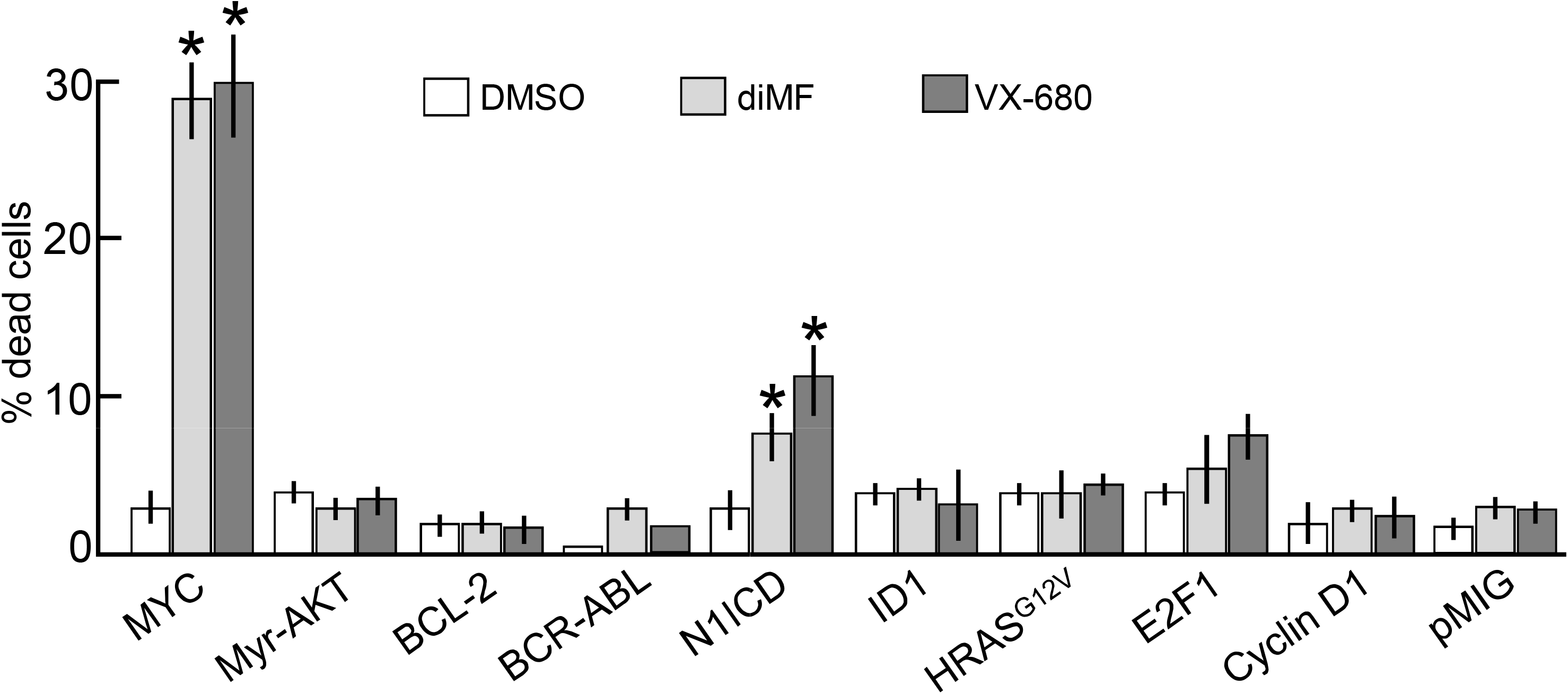
diMF is not synthetic lethal with a variety of other oncoproteins. Rat1A cells harboring either an empty vector or one of the indicated oncogenes, were treated with either 0.1% DMSO (□), 6 μM diMF (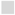) or 300 nM VX-680 (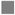) for 3 days. Cell viability, was assayed using the Trypan blue exclusion assay. Each data point is the average of three independent experiments, with treatments done in triplicate (n=9). Error bars denote standard deviation. For cells overexpressing MYC and the N1ICD, viability of the diMF and VX-680 groups was statistically different from DMSO (one-way ANOVA followed by Dunnett’s test). The symbol * indicates *p<0.01* for drug treated compared to DMSO control.

**Figure 6:**
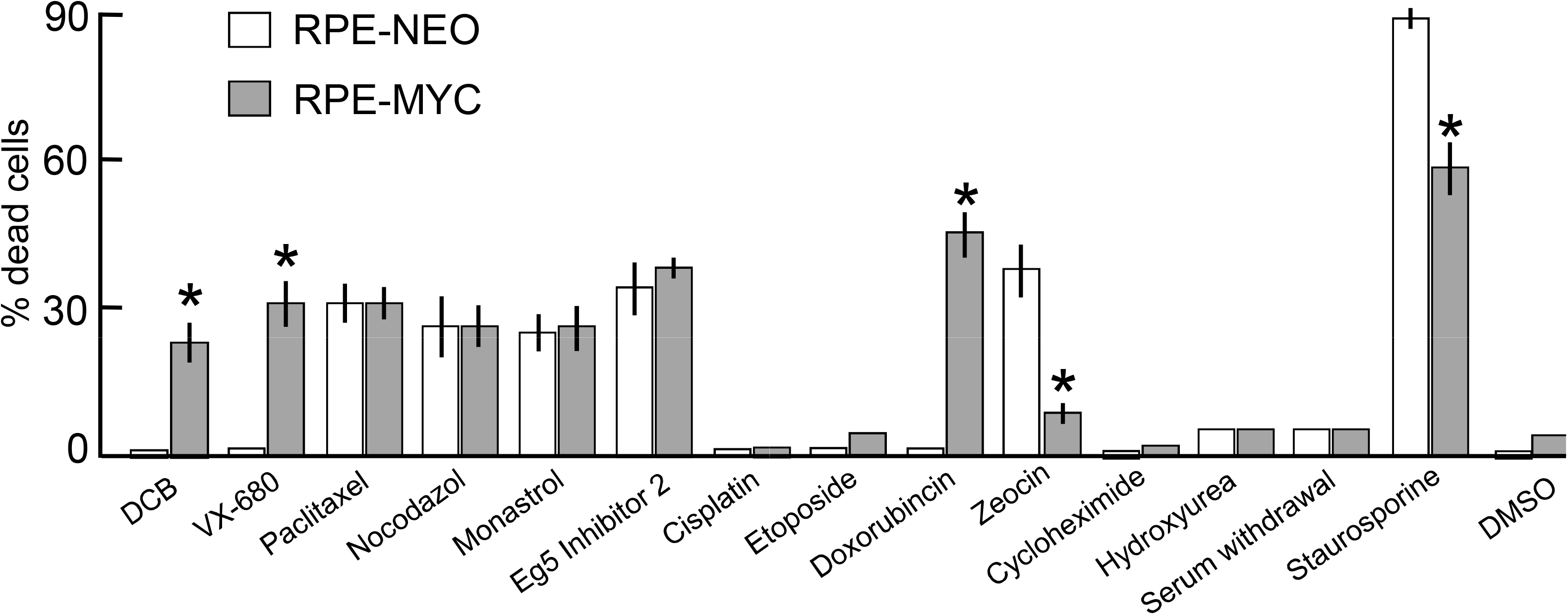
MYC-diMF synthetic lethality is not due to a general priming of apoptosis. Cells overexpressing (RPE-MYC) or not overexpressing (RPE-NEO) MYC were treated for 72 hours with various chemical primers of apoptosis: DCB (2μg/ml), VX-680 (300 nM), Paclitaxel (5 μM), Nocodazole (1μg/ml), Eg5 inhibitor II (10μM), Monastrol (20 μM), Doxorubicin (200 nM), Cisplatin (200 μM), Etoposide (20 μM), Zeocin (50 μg/ml), Cycloheximide (10 μg/ml), Staurosporine (10 μM), serum withdrawal and 0.1% DMSO as a control group. Cell viability was assayed by Trypan blue exclusion assay. Each column represents the average of three independent experiments, with three replicates in each experiment (n=9). Error bars denote standard deviation. Viability differences between RPE-NEO and RPE-MYC cells with each chemical were statistically compared using two tailed, unpaired t-tests. The symbol * indicates *p<0.01*.

## Results

### Selection of the ROCK inhibitor diMF in a screen for synthetic lethality with MYC

MYC is known to induce potent pro-apoptotic activity but only under specific conditions of cellular stress [1, 19, 20]. We assayed more than 2700 compounds for activity that recapitulated the cellular contexts under which cells could not survive with high levels of the MYC oncogene. Previously, we generated a pair of isogenic cell lines, RPE-NEO and RPE-MYC, by engineering human retinal pigment epithelial (RPE) cells to ectopically express a neomycin resistance gene and the *MYC* oncogene, respectively. This pair of cell lines has been used to demonstrate synthetic lethal interactions between overexpression of MYC and inhibition of both CDK1 [13] and AURKB [15].

Previously, synthetic lethality with the Aurora kinase inhibitor VX-680 has been attributed to the ability of the compound to elicit a transient mitotic arrest and then block cytokinesis [15]. This results initially in apoptosis, but also the eventual accumulation of polyploid cells. Our screen aimed to identify chemicals that can elicit similar cell-division defects. To facilitate screening, a histone *H2B-GFP* fusion gene was stably transfected into cells. This enabled imaging of DNA in live cells. Events were considered positive for synthetic lethality with MYC when cell death was present in RPE-MYC cells, but absent in RPE-NEO cells.

One compound that elicited extensive mitotic arrest at 24 hours and polyploidy at 48 hours (Fig. 1A) was identified as dimethylfasudil (diMF), (*S*)-(+)-2-Methyl-1-[(4-methyl-5-isoquinolinyl)sulfonyl]-hexahydro-1*H*-1,4-diazepine dihydrochloride, a competitive inhibitor of ROCK kinases (Fig. 1B). With an IC_50_ of 12 nM for ROCKII inhibition, diMF is considered a potent, selective and reversible ROCK inhibitor. It has only been used in preclinical *in vitro* and *in vivo* studies [21], perhaps due to the plethora of alternative ROCK inhibitors that have been developed [22]. In our confirmatory assays, diMF killed 30% of RPE-MYC cells within an initial 3-day treatment, whereas RPE-NEO did not succumb (Fig. 1C). The solvent for diMF (DMSO) failed to elicit cell death in either RPE-NEO or RPE-MYC cells (Fig. 1C). An important observation from our screen is that fasudil and ripasudil, both potent ROCK inhibitors, did not elicit cell death or polyploidy. Thus, diMF alone among the ROCK inhibitors in the screen, could selectively kill cells that overexpress MYC.

### diMF-induced death in MYC overexpressing cells is mediated by a mitochondrial apoptotic pathway

To investigate the nature of MYC-dependent death elicited by diMF, we examined the features of diMF-induced cell death in RPE-MYC cells. At least a portion of the cell death phenotype was inhibited by the pan-caspase inhibitor z-VAD-fmk. At 72 hours after 6 μM diMF treatment, z-VAD-fmk at 100 μM provided more than 50% protection against cell death (Fig. 1C). Morphologically, we observed early classical features of apoptosis after diMF treatment such as shrinkage of the cell and condensation and fractionation of nuclear chromatin into sharply delineated regions (Fig. 2A). Cleaved, or activated, forms of caspases 9 (Fig. 2B, C) and 3 (Fig. 2C) and the downstream caspase target poly ADP-ribose polymerase 1 (PARP1) were found after diMF treatment (Fig. 2C). These features were likely due to activation of mitochondria-dependent apoptosis, since we also could demonstrate diMF-dependent dissipation of the mitochondrial membrane potential, ΔΨm, in RPE-MYC cells (Fig. 2C). These observations support the conclusion that at least some of the MYC-dependent, diMF-induced cell death is through a mitochondrial apoptotic cascade.

### Acute activation of MYCN also confers susceptibility to the cytotoxicity of diMF

Arguably, the model RPE cell lines used in our experiments have been propagated extensively and it is possible that sustained expression of MYC selected for genetic or epigenetic alterations that were responsible for the observed interaction with diMF, not MYC itself. To dispel this possibility and also to determine if another MYC paralog could substitute for MYC in the synthetic lethality, we examined sensitivity to diMF in cells expressing MYC^ER^ and MYCN^ER^ fusion proteins, where MYC and MYCN are fused with the estrogen receptor. In these cells, MYC activity is acutely activated by the addition of 4-hydroxy-tamoxifen (4-OH) which induces translocation of oncogenic MYC fusion protein to the nucleus [4]. While both 4-OH and diMF themselves resulted in low but statistically insignificant levels of cell death, the drugs acted synergistically to elicit approximately 30% overall lethality (Fig. 3A-D). This was true with MYC^ER^ in both HA1E (human kidney epithelial cells) and IMR90 (human lung fibroblast) cells. It was also apparent in MYCN^ER^ expressing Rat1a (rat embryo fibroblast) and HA1E cells. We conclude that overexpression of either MYC or MYCN is lethal in combination with diMF, and that the synthetic lethal relationship is conserved in rodent and human cells and both fibroblast and epithelial cell lineages.

### Depletion of MYC negates sensitivity diMF

Human cancers frequently overexpress MYC, but simultaneously may harbor additional mutations, copy number alterations and translocations. The products of these additional genetic events may mask the synthetic lethality we observed with model immortalized cell lines. We assayed if human cancer-derived cells with abundant MYC were sensitive to diMF and whether overexpression of MYC was necessary for sensitivity. The human cancer lines Hela (cervical), Calu-6 (lung adenocarcinoma), NCI-H841 (small cell lung cancer) and DU-145 (prostate) are all known to express MYC abundantly and treatment with diMF for four days killed 25-35% of cells with all four of these cell lines (Fig. 4A). Next, to assay if MYC was absolutely required for diMF sensitivity, we used RNAi to deplete MYC. MYC protein levels were markedly reduced by MYC RNAi (estimated reduction to < 20% control) (Fig. 4B). Depletion of MYC alone elicited no appreciable levels of toxicity in any of the cell lines (Fig. 4C). RNAi depletion of MYC did elicit a more flattened morphology, a change previously reported [18], but cells remained viable. However, depletion of MYC with RNAi did provide each cell line with a 50-75% protection from diMF-induced cytotoxicity (Fig. 4D). These findings indicate that abundant MYC, although dispensable for survival in the absence of diMF, was required for cells to undergo apoptosis in response to diMF.

### Alternative oncogenic events do not substitute for MYC in diMF-induced lethality

Our observation with MYC knockdown prompted us to examine more closely if other oncogenic proteins, especially those that influence the cell cycle, might substitute for MYC and confer synthetic lethality with diMF. We assayed the possibility using a panel of rat embryo fibroblasts (Rat1A) that have been engineered to ectopically express a variety of cancer genes [13]. The oncogenic proteins expressed included oncogenic HRAS^G12V^, myristoylated AKT1 (MyrAKT), BCL-2, ID-1, E2F1, the intracellular domain of NOTCH1 (N1ICD), CYCLIN D1, and BCR-ABL [13]. In a blinded assay, only MYC and the N1ICD-expressing Rat1a cells demonstrated statistically significant cell death with 72 hours of diMF treatment (Fig. 5). We concluded that diMF elicited synthetic lethality in combination with MYC and, with the possible exception of the N1ICD, its role could not be substituted by the other oncoproteins assayed. However, N1ICD-diMF lethality was only a fraction of that seen with elevated MYC and diMF treatment and NOTCH1 is a known activator of MYC transcription [23–25]. This regulation may not be conserved in Rat1a cells [16], so whether or not the N1ICD-diMF interaction is a recapitulation of MYC-diMF synthetic lethality has yet to be definitively determined.

Our findings are similar to data obtained in RPE-MYC cells with the pan-Aurora kinase inhibitor VX-680 or an AURKB-specific inhibitor AZD1152 [15]. We recapitulated the findings with VX-680 in the Rat1a panel (Fig. 5). Ultimately, synthetic lethality with VX-680 has been attributed to disabling the chromosomal passenger protein complex (CPPC) [15]. Our findings suggest that like VX-680, diMF might somehow disable mitotic spindle assembly, the CPPC or other complexes essential to mitosis, a role not previously ascribed to diMF.

### Overexpression of MYC sensitizes cells only to selective apoptotic stimuli

We next asked whether any drug-mediated disturbance of cell division could selectively kill cells that overexpress MYC. We utilized chemical inhibitors to explore four additional means to disturb cell division: (1) stabilizing mitotic spindle microtubules from disassembly with paclitaxel, (2) interfering with the polymerization of microtubules with nocodazole, (3) targeting the centrosomal motor protein Eg5 (also known as KIF11) with monastrol [26], and (4) interfering with assembly of the cytokinesis contractile-ring microfilaments with dihydrocytochalasin B (DCB).

Prevention of cytokinesis with DCB mimicked the selective killing of diMF and VX-680 specifically in RPE-MYC cells (Fig. 6). In contrast, paclitaxel and nocodazole killed both RPE-NEO and RPE-MYC. Likewise, inhibition of the centrosomal motor protein Eg5 with either monastrol or Eg5 inhibitor II, both of which prevent separation of centrosomes to form the bipolar mitotic spindle [26], killed cells without discrimination. Both the spindle toxins and Eg5 inhibitors caused prolonged arrest in mitosis prior to apoptotic cell death. In contrast, prolonged arrest in mitosis was not associated with treatment with diMF, DCB and VX-680. It may be that apoptosis in RPE cells following a prolonged arrest in mitosis is not primed by overexpression of MYC, but this is speculative.

A variety of other apoptotic stimuli failed to sensitize RPE-MYC cells to apoptosis. For example, the deoxyribonucleotide synthesis inhibitor hydroxyurea, protein synthesis inhibitor cycloheximide, and serum deprivation all arrested cellular proliferation but caused no cell death in both RPE-NEO and RPE-MYC. Overexpression of MYC sensitized cells to doxorubicin, yet cisplatin and etoposide elicited proliferative arrest, rather than cell death, in both cell lines. Zeocin effectively kills RPE-NEO cells, but not RPE-MYC cells.

Collectively, these data imply that overexpression of MYC does not always prime cells to apoptosis, even with drugs that directly target the mitotic spindle or centrosomes. Doxorubicin which intercalates with DNA and inhibits topoisomerase II during DNA replication was preferentially lethal in RPE-MYC cells, but polyploid cells were not induced by this drug and they were induced with diMF (Fig. 1B). Overall, the data points to a target for diMF that is active at or upstream of the CPPC or functions during cytokinesis.

## Discussion

MYC contributes to the genesis of many cancers and although it has been a focus of drug research, strategies to target or exploit MYC have yet to be clinically successful. Direct targeting of MYC with small molecule inhibitors may, in fact, be technically impractical. Alternatively, synthetic lethal approaches provide promise to selectively kill tumor cells that overexpress MYC while sparing normal somatic cells [27].

One limitation with current MYC synthetic lethal compounds is toxicity. To circumvent this, we attempted to repurpose existing small-molecule drugs. We carried out phenotypic screening of a small molecule library for MYC-synthetic lethal agents. The well characterized MYC synthetic lethal phenotype elicited by Aurora kinase B inhibitors [15, 16, 28, 29] served as a measure for our screen. MYC-VX-680 synthetic lethality, for example, induces transient mitotic arrest and apoptosis, followed by induction of polyploidy [15]. The screen uncovered similar activity with a Rho kinase inhibitor, diMF. Our confirmatory experiments showed that this agent selectively elicits lethality with MYC activity. A variety of oncogenic manipulations failed to substitute for MYC in sensitizing cells to diMF. The MYC-diMF synthetic lethality is therefore not a consequence of cellular transformation. In addition, a variety of apoptosis-inducing agents did not preferentially kill MYC-expressing cells, so a general priming of apoptotic signaling by diMF was not responsible for the synthetic lethality either. Instead, diMF likely targets a unique vulnerability induced by MYC overexpression.

ROCK 1 and ROCK 2 belong to the AGC (PKA/PKG/PKC) family of serine-threonine kinases [22]. diMF has been extensively studied as a ROCK inhibitor and characterized in pre-clinical models. It promotes generation of pancreatic beta-like cells from human pluripotent stem cells [30], augments neurite extension [31], enhances proliferation and migration of endothelial progenitor cells [32], and triggers cell-cycle arrest and cellular senescence of mouse embryo fibroblasts [33]. However, ROCK-independent activity has also been ascribed to diMF. Degradation of polyglutamine-expanded ataxin-3 and 7, causative in spinocerebellar ataxias, is induced by diMF independent of ROCK inhibition [34]. Others have reported non-canonical targets of diMF include LRRK2 [35] and PKA [36]. The MYC synthetic lethality observed here also appears to be ROCK-independent. Other ROCK kinase inhibitors in the screening library, including fasudil and ripasudil, were not identified by our screening effort and later were confirmed negative in eliciting apoptosis and polyploidy in RPE-MYC cells (data not shown).

One report links diMF to mitotic defect and polyploidy. Polyploidy and apoptosis of malignant megakaryocytes can be induced by diMF and this has been ascribed to the inhibition of AURKA [37]. In RPE-MYC cells, MYC synthetic lethality is induced by the AURKB-specific inhibitor AZD-1152 [15]. Therefore, diMF may target multiple AURK members or unknown CPPC components. Nonetheless, we show here that diMF only elicits apoptosis and polyploidy in cells that overexpress MYC. Elucidating diMF targets definitively responsible for this synthetic lethality will require further research.

Fasudil and ripasudil are used clinically and have close structural similarity to diMF. We believe this means diMF will also have drug-like properties; the favorable activity, pharmacokinetic and toxicity profile required for clinical applications. Our findings raise the possibility that diMF might be repurposed for the treatment of MYC overexpressing tumors, which are generally aggressive and may currently lack a targeted therapy option.

## Acknowledgments

We thank Andrei Gaga (University of California, San Francisco) for the Rat1A panel of cells and the H2G-GFP expression construct, Mariia Yuneva (The Francis Crick Institute) for HA1E/Myc^ER^, HA1E/MycN^ER^ and IMR90/Myc^ER^ cell lines and Sue Kim (University of Arizona) for the Rat1A/MycN^ER^ cell line. This research was supported by funds from J. Michael Bishop Institute of Cancer Research (to J.Z. and D.Y.).

